# Skin stem cells orchestrate *de novo* generation of extrathymic regulatory T cells to establish a temporary protective niche during wound healing

**DOI:** 10.1101/2021.08.16.456570

**Authors:** Cynthia Truong, Weijie Guo, Liberty Woodside, Audrey Gang, Peter Savage, Nicole Infarinato, Katherine Stewart, Lisa Polak, John Levorse, Amalia Pasolli, Stanislav Dikiy, Alexander Rudensky, Elaine Fuchs, Yuxuan Miao

**Affiliations:** Robin Neustein Laboratory of Mammalian Development and Cell Biology, Howard Hughes Medical Institute, The Rockefeller University, New York, NY 10065, USA; Ben May Department of Cancer Research, The University of Chicago, Chicago, IL, 60615, USA; Department of Pathology, The University of Chicago, Chicago, IL, 60615, USA; Electron Microscopy Resource Center, The Rockefeller University, New York, NY 10065, USA; Immunology Program, Sloan Kettering Institute, Howard Hughes Medical Institute, Memorial Sloan Kettering Cancer Center, New York, NY 10065, USA

## Abstract

Adult stem cells reside in various tissues to govern homeostasis and repair damage. During wound healing, these stem cells must be mobilized to enter the center of the injury where they are exposed to many inflammatory immune cells infiltrating the wounded tissue. While these immune cells are indispensable for preventing infections and clearing dead cells, they can also create a harsh inflammatory environment which could potentially damage the stem cells and prevent their self-renewal and differentiation. Here, using a model of cutaneous wound healing in which hair follicle stem cells (HFSCs) repair the wound, we show that, upon migrating into the wound, skin stem cells acquire a strong immune modulatory capacity which allows them to sculpt a temporary immune suppressive niche for self-protection. We reveal that the HFSCs in the wound bed orchestrate extrathymic differentiation of regulatory T (Treg) cells by providing co-stimulation to the wound-infiltrating CD4 effector T cells. In this way, Treg cells can be generated *de novo* in close proximity to and can intimately protect HFSCs from the collateral damage inflicted by inflammatory neutrophils. This study uncovered a striking inflammatory adaptation capacity unique to adult tissue stem cells which allows them to shape their own immune suppressive niche during wound repair.

## Introduction

Adult stem cells (SCs) are responsible for the maintenance and regeneration of various tissues and appendages (Almada and Wagers, 2016; Blanpain and Fuchs, 2009; Goodell et al., 2015). As a group of indispensable, long-lived cells, they are challenged over their life cycles by frequent exposure to waves of inflammation caused by infection or injury(Naik et al., 2018; Niec et al., 2021). In order to ensure tissue homeostasis while maintain the SC pool, SCs must be protected from these repeated bouts of inflammation. Despite the physiological significance of this feature, it is unclear how adult SCs achieve self-renewal and differentiation within an inflammatory context.

Mouse skin offers an excellent system to tackle these questions. In the skin, the hair follicle stem cells (HFSCs) reside within the bulge, isthmus, and infundibulum regions of the hair follicle, located in the dermis. These stem cells not only fuel hair regeneration, but they also contribute to the repair of damaged epidermis. Conventionally, it is thought that these SCs evade damage from inflammation by residing within an immune-privileged niche (Agudo, 2020; Ichiryu and Fairchild, 2013). The hair follicle niche may maintain immune privilege by: firstly, providing a physical barrier to separate HFSCs from any infiltrating inflammatory cells, and secondly, by producing immunosuppressive molecules such as IL-10 or TGFβ to dampen inflammation around the HFSCs. Additionally, suppressive immune cells, such as regulatory T cells (Treg) are deployed near the hair follicle and can also potentially protect the HFSCs(Ali et al., 2017).

These niche factors may be able to provide adult SCs with a protective barrier against inflammation in homeostasis. During wound healing, however, SCs must be mobilized to leave their niche and migrate into the wound bed to restore the barrier (Gonzales and Fuchs, 2017). While this is ongoing, the damaged tissue is exposed to pathogens and dead cells, triggering a cascade of responses which includes the recruitment of inflammatory immune cells such as neutrophils, monocytes, and macrophages, which then secrete a variety of anti-microbial molecules as well as pro-inflammatory cytokines. These factors are critical for fending off pathogens and dead cells, but if left unchecked, can cause significant collateral stress for the SCs. Caught in the crosshairs of such robust inflammation, and outside of their immune-privileged niche, it is unknown how adult SCs are still able to evade collateral damage from infiltrating immune cells in order to successfully re-epithelialize the wound.

Interestingly, recent studies on tumor-initiating stem cells have revealed that these tumor SCs are endowed with unique molecular features to resist robust anti-tumor immunity (Miao et al., 2019). These tumor SCs achieve this effect through directly modulating the activities of antitumor immune cells (Miao et al., 2019). These observations strengthen the speculation that adult SCs are equipped with unique programs to adapt to inflammation. Since wounded and tumorigenic stem cells share many molecular features(Ge et al., 2017), it is possible that the mechanisms utilized by tumor SCs to evade immune attack are co-opted from normal stem cell programs intended to shield SCs from inflammation during tissue repair.

In this study, we employed a mouse model of partial-thickness removal cutaneous wounding, in which HFSCs are known to become activated and migrate out of their niche and into the highly inflammatory wound bed to re-epithelialize the wound. We unveil that after migration, HFSCs are able to activate a strong immune modulatory program which is capable of inducing *de novo* generation of Treg cells. Importantly, the intimate crosstalk between HFSCs and Treg cells functions to actively sculpt a temporary immune suppressive niche which prevents accumulation of inflammatory neutrophils around the wound-repairing HFSCs.

## Results

### Hair follicle stem cells activate immune modulatory programs during wound repair

Partial-thickness wounding, which mechanically removes the superficial epidermis while leaving the dermal components intact, mobilizes HFSCs to migrate out of their niche and into the wound bed to repair the upper hair follicle and epidermis (Ge et al., 2017) (Figure 1A). This wounding model provides an ideal system to dissect the mechanisms underlying adult SCs’ ability to adapt to inflammation outside of their immune privileged niche. By using *Sox9CreER;Rosa26-tdTomato* mice to trace the behavior and fate of HFSCs, we confirmed that, by the third day after wounding, HFSCs have begun to migrate out of their hair follicle niche and have entered the wound bed (Figure 1A). By day five, the new epithelium is regenerated and comprised primarily of the progeny of these migratory HFSCs (Figure 1A). That these migratory HFSCs can successfully regenerate epidermis within an inflammatory environment but outside of their original niche suggests that these HFSCs are able to successfully adapt to inflammation by themselves. To identify the molecular programs employed by HFSCs to protect themselves from inflammation in the wound, we used fluorescence-activated cell sorting (FACS) to isolate Tomato+ HFSCs from unperturbed skin, as well as from wounded skin at day 3 post-wounding, when the HFSCs have migrated to the wound bed (Supplementary Figure 1A). We then subjected these cells to single cell RNA-sequencing. GO term analysis revealed that genes involved in regulation of immune responses are significantly enriched in HFSCs upon wounding (Supplementary Figure 1B). Surprisingly, *Cd80* was found to be activated in the wounded HFSCs (Figure 1B). CD80 is a cell-surface ligand canonically thought to be specifically expressed by immune cells, in particular by professional antigen presenting cells such as dendritic cells or macrophages. In this context its function is to modulate immune cell responses by providing co-stimulatory signals to T lymphocytes. CD80 has recently been found to be selectively acquired by tumor-initiating stem cells in squamous cell carcinomas, where it functions to directly suppress cytotoxic T cell-mediated anti-tumor immunity. (Miao et al., 2019). Analysis of the chromatin accessibility of HFSCs before and after wounding using Assay for Transposase-Accessible Chromatin using Sequencing (ATAC-seq) confirmed that the *Cd80* promoter gained a strong peak upon wounding (Figure 1C). Our single cell analysis further confirmed that *Cd80* can be activated on CD45-non-hematopoietic SCs (Supplementary Figure 1C). Interestingly, *Cd80* was found to be specifically activated on the HFSCs that had acquired Integrin a5 (Figure 1B), undergone epithelial-mesenchymal transition, and acquired migratory capacity (Supplementary Figure 1D). This restricted expression of CD80 was also confirmed by flow cytometry, Imagestream, and Immunofluorescent staining, which showed specific upregulation of CD80 on migratory HFSCs that have exited the hair follicle and entered the wound (Figure 1D, E, F). Intriguingly, HFSCs don’t appear to activate *Cd86*, the other costimulatory ligand thought to be redundant with CD80 on immune cells (Supplementary Figure 2A). Therefore, the unique expression pattern of CD80 implicates that this immune modulatory ligand might play an important role for HFSCs outside of their natural niche during wound repair.

**Figure 1.**
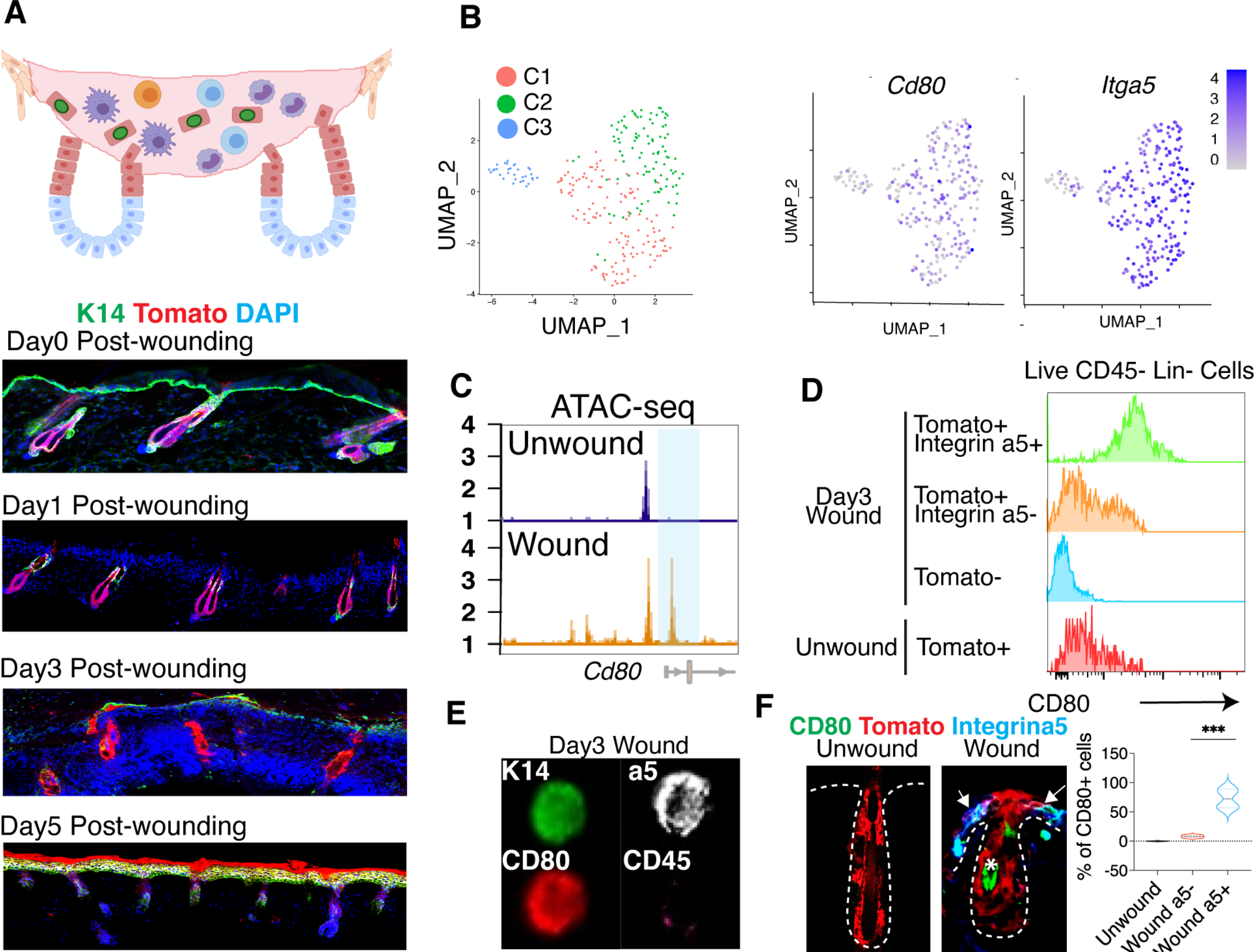
HFSCs activate CD80 upon migrating into the wound. **(A)** Schematic and immunofluorescent images showing the healing dynamics of partial thickness removal cutaneous wound by HFSCs **(B)** MA plot displaying RNA-seq analysis of HFSCs isolated from unwounded versus wounded skin **(C)** t-SNE plots showing *Cd80* expression on wounded Integrin a5-expressing HFSCs **(D)** ATAC-seq peaks showing *Cd80* activation upon wounding **(E-F)** Flow cytometry (E), Imagestream (F) and Immunofluorescence staining/quantification confirming CD80 activation on CD45-Integrin a5+ HFSCs in wounded skin.

### CD80 is important for HFSCs to repair cutaneous wounds

To test the functional significance of CD80 for HFSCs, we first took advantage of the finding that HFSCs are the major non-hematopoietic cells that express CD80 in wounded skin (Figure 1D). Based on this result, we reconstituted lethally irradiated CD80-null mice with wild-type bone marrow and generated bone marrow chimeras (CD80 BMC). In these mice, CD80 expression was reconstituted in immune cells, and remained lost only in HFSCs during wounding. As a control, we irradiated WT mice and reconstituted them with WT BM (WT BMC). By using the congenic marker CD45.1 to distinguish donor cells from recipient endogenous immune cells, we confirmed that in the wound, the majority of myeloid cells, which are the major population supplying immune cell-derived CD80-mediated co-stimulation, are derived from CD80-competent WT bone marrow (Supplementary Figure 2B). At day 1 post-wound, in both CD80 BMC and control BMC mice, the superficial epidermal basal keratinocyte layer (K14+) is stripped, with no gross phenotypic differences between CD80 KO and control. In the dermis, the hair follicle bulges remain intact (Figure 2A). At day 3 post-wound, the re-epithelization process is initiated in both control and CD80 BMC at a similar rate (Figure 2A). Strikingly, by day 5, whereas the wound in control WT BMC mice is completely re-epithelialized with hyperproliferative K14+ cells spanning the length of the wound, the CD80 BMC mice display only discrete portions of re-epithelialization, with large areas consisting of only granulation tissue (Figure 2A). Additionally, the differentiation program of HFSCs is significantly disrupted in CD80 BMC mice. While extensive K10+ suprabasal cells are generated in control BMC mice by day 5 post-wound, the number of differentiated cells is drastically reduced in CD80 BMC mice (Figure 2A).

**Figure 2.**
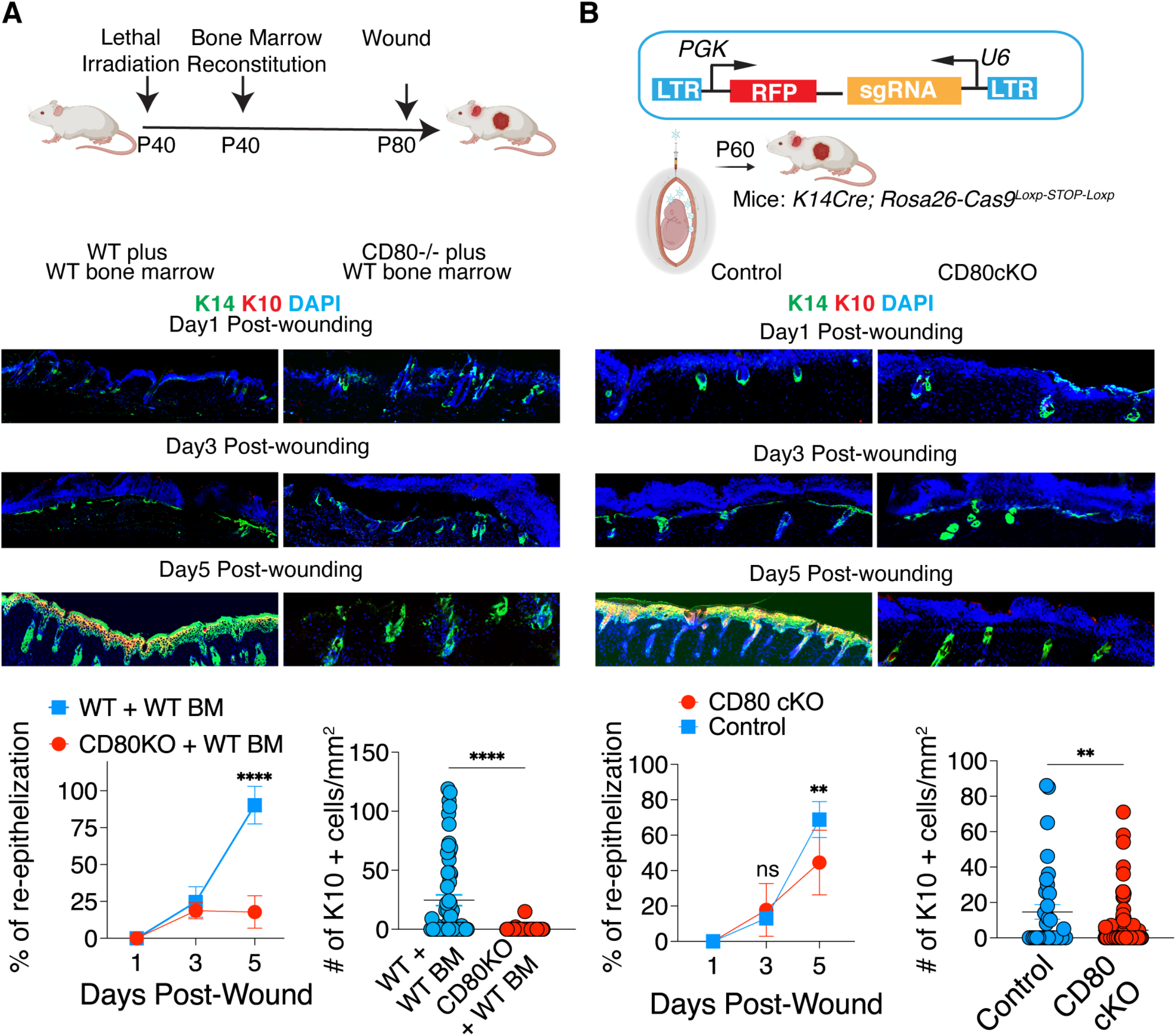
HFSCs activate CD80 upon migrating into the wound. **(A)** IF staining and quantification of percentage of wound re-epithelialization (K14+) and number of differentiated cells (K10+) in wounded CD80 bone marrow chimera mice, demonstrating the importance of CD80 for wound healing **(B)** IF staining and quantification of percentage of wound re-epithelialization (K14+) and number of differentiated cells (K10+) using *in utero* injection of a Cd80 sgRNA-containing lentiviral construct to achieve CD80 cKO, demonstrating the importance of skin-specific CD80 for wound healing

Next, we sought to design additional models to ensure that CD80 is silenced only in skin and to further interrogate its functional significance for wound repair. For this purpose, we used a powerful ultrasound-guided *in utero* lentiviral (LV) microinjection approach to transduce the single-layered surface of mouse ectoderm at E9.5 of mouse embryonic development. This technique enables selective delivery of sophisticated genetic constructs into the skin progenitors, allowing us to achieve rapid genetic manipulation of the skin (Beronja et al., 2010). With this approach, we injected a lentiviral construct containing a small guide RNA (sgRNA) targeting *Cd80* into the skin progenitors of *K14Cre;Rosa26-Cas9 ^Loxp-STOP-Loxp^* embryos. In these mice, Cre expression is driven by the promoter specific for the basal epithelium, such that CRISPR/Cas9-mediated gene editing is activated only in the basal layer of keratinocytes where the HFSCs are located. In our model, this induced conditional CD80 silencing (CD80 cKO) specifically in the skin (Supplementary Figure 2C). Upon wounding, while Cre-negative control mice completed re-epithelization and differentiation by day 5 post-wounding, both the re-epithelialization and differentiation processes were significantly disrupted in CD80 cKO mice (Figure 2B).

### SC-CD80 orchestrates Treg responses to prevent accumulation of neutrophils near migratory HFSCs

Intrigued by the striking impact of CD80 on the wound healing capacity of HFSCs, we examined the underlying mechanisms that may allow stem cell CD80 to promote wound repair. CD80 is canonically expressed on antigen presenting cells, where it engages with CD28 to provide co-stimulatory signals to T lymphocytes (Belkaid and Oldenhove, 2008). The presence of this marker on migratory HFSCs during wounding hinted at the tantalizing possibility that HFSC-CD80 may function to shape the immune environment of HFSCs during wounding. To test this hypothesis, we profiled the immune cell compositions of our mice by flow cytometry. In the unwounded state, both CD80 BMC or CD80 cKO mice and control skin exhibited no differences in immune cell composition (Supplementary Figure 2D). Interestingly, significantly more immune cells (CD45+) infiltrated the wound in the CD80 cKO mice, (Figure 3A), suggesting overwhelming inflammation. However, we observed a significant deficit in both the percentage and absolute numbers of CD4 T cells (Figure 3B), particularly the CD4+Foxp3+ Regulatory T cells (Treg cells) in both CD80 BMC and CD80 cKO mice compared to control mice at day 5 post-wound (Figure 3C). Treg cells are a key group of suppressive immune cells, preventing deleterious autoimmunity and dampening down inflammation (Josefowicz et al., 2012; Sakaguchi et al., 2020). During cutaneous wound repair, depletion of Tregs has been shown to attenuate wound closure (Mathur et al., 2019; Nosbaum et al., 2016). Consistent with previous studies, we wounded Foxp3-DTR mice, which allowed for selective and inducible depletion of Tregs in the skin upon administration of diphtheria toxin (DT), and found that these mice, similar to our CD80 BMC and CD80 cKO models, were defective in re-epithelialization and differentiation (Figure 3D). These data support the conclusion that HFSCs may promote a robust Treg response to support wound healing.

**Figure 3.**
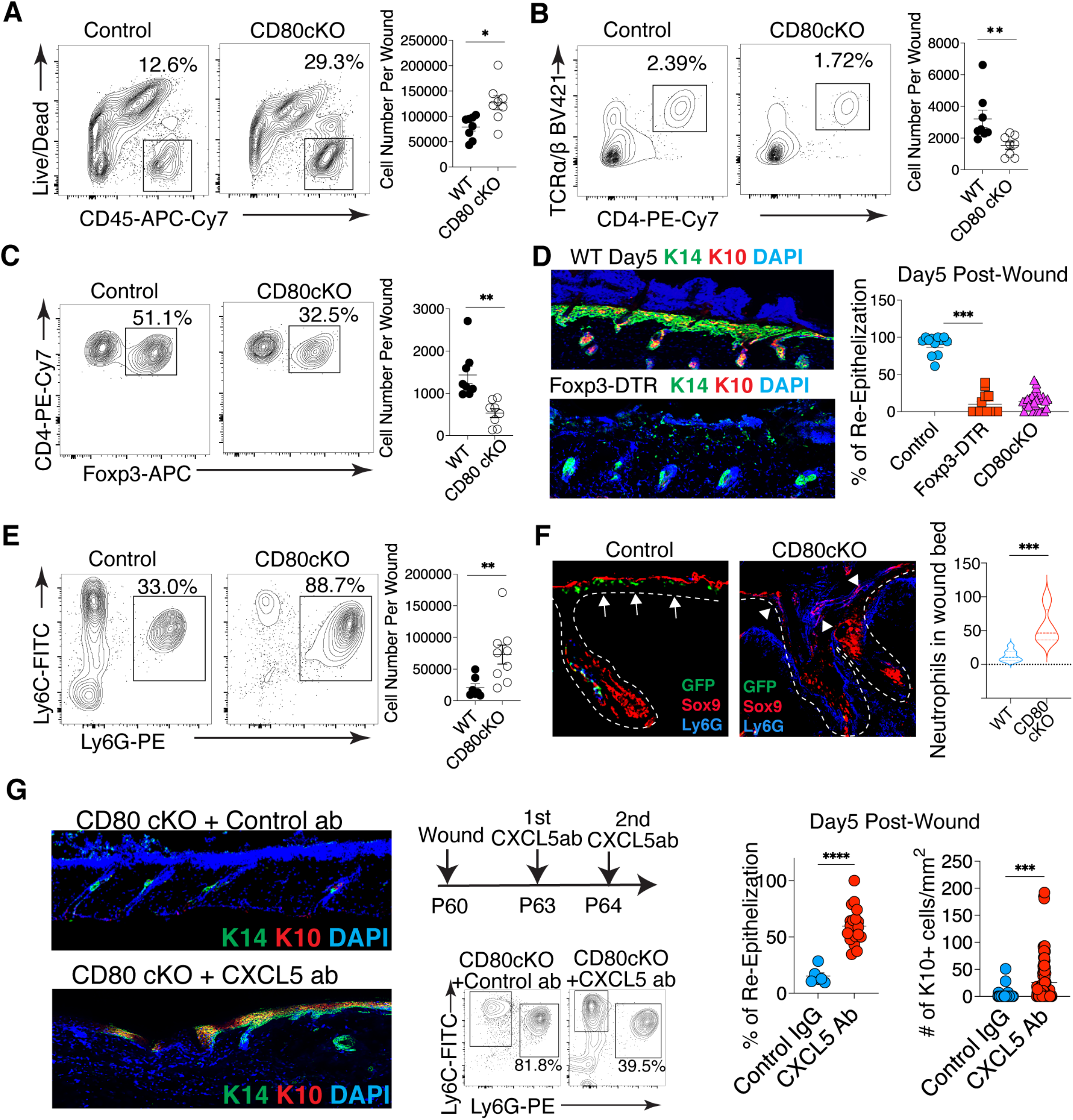
CD80 on HFSCs is critical for orchestrating Treg-mediated suppression of neutrophil response during wound healing. **(A-C)** Flow cytometry quantification showing an increase in total CD45+ immune cells (A) but a decrease in CD4+ T cells (B), in particular CD4+ Foxp3+ Tregs (C), in CD80 cKO wounds. **(D)** IF staining and quantification of percentage of re-epithelialization (K14+) and number of differentiated cells (K10+) in Treg-depleted mice, demonstrating the importance of Tregs for wound healing. **(E)** Flow cytometry quantification showing the accumulation of CD11b^+^MHCII^-^Ly6G^Hi^ Ly6C^low^ neutrophils in the CD80 cKO wound. **(F)** IF staining and quantification showing the accumulation of Tregs within the wound bed in close proximity to Sox9+HFSCs, preventing neutrophils from approaching Sox9+ cells in WT wounds (left). In contrast, in CD80 cKO wounds, Tregs are absent and neutrophils accumulate in the wound bed (right). **(G)** IF staining and quantification of percentage of re-epithelialization (K14+) and number of differentiated cells (K10+) in neutrophil-depleted mice demonstrating the neutrophil-mediated delay in wound healing seen in CD80 cKO mice.

Given the vital role of HFSC-Treg dialogues during wound repair, we were curious about which immune populations were controlled by this crosstalk. Immune profiling showed that in CD80 cKO mice, the reduction of Treg cells was accompanied by a drastic increase in neutrophils (Figure 3E). Strikingly, immunofluorescence revealed that in WT mice, Treg cells accumulate in the wound bed and are localized in close proximity to migratory HFSCs, separating the HFSCs from the neutrophils (Figure 3F). However, in CD80 cKO mice, Treg cells are rarely detected next to migratory HFSCs, and as a result, neutrophils extensively infiltrated and accumulated in the wound bed and hair follicles underneath the wound (Figure 3F). Recent studies have indeed shown that the Treg cells can indirectly reduce neutrophil infiltration by suppressing CXCL5 production (Mathur et al., 2019). To test whether excess neutrophil accumulation in the wound bed was the root cause of impaired re-epithelialization upon loss of HFSC-CD80, we treated CD80cKO mice with an anti-CXCL5 antibody prior to wounding (Mathur et al., 2019). Neutrophil depletion significantly rescued the re-epithelialization and differentiation defects seen in CD80 cKO mice (Figure 3G). These data provide compelling evidence that HFSC-CD80 plays a critical role in orchestrating Treg responses, which in turn prevent excess neutrophil accumulation during wound healing.

### HFSCs facilitate extrathymic *de novo* generation of Treg cells in the wound

We next sought to investigate the mechanism by which HFSCs modulate Treg responses through CD80. It is well established that Treg cells develop either ontogenetically in the thymus (tTreg) or inducibly in the periphery (iTreg). The B7 molecules expressed by antigen presenting cells have been shown to increase proliferation of Treg cells (Salomon et al., 2000). We designed experiments to distinguish the two equal possibilities: that either HFSCs were promoting the proliferation/expansion of recruited tTregs, or that they were inducing the *de novo* generation of iTregs within the wound. To this end, we first measured the proliferation of Treg cells in WT and CD80 cKO mice. Intriguingly, we did not detect any significant difference in the Treg cell Ki67 level between WT and CD80 cKO mice (Figure 4A). Next, we designed a Treg fate-mapping assay in which preexisting tTreg cells could be distinguished from any extrathymically-induced Treg cells generated during wound healing. To this end, *Cd80^-/-^* versus WT mice were reconstituted with bone marrow extracted from *FoxP3GFP-CreER;Rosa26-tdTomato^Loxp-STOP-loxp^* mice. In these chimeric mice, a fate-mapping marker activated by tamoxifen prior to wounding marked all pre-existing Treg cells with both tdTomato and GFP. In contrast, any new iTreg cells generated *de novo* during wounding were marked solely by GFP, and not with Tomato (Figure 4B). Interestingly, wounding significantly increased the percentage of GFP single positive Treg cells in WT wounds, but not in CD80 KO wounds. This result strongly suggests that HFSC-CD80 is critical for inducing *de novo* generation of new Tregs, rather than promoting the expansion of existing Treg cells during wounding.

**Figure 4.**
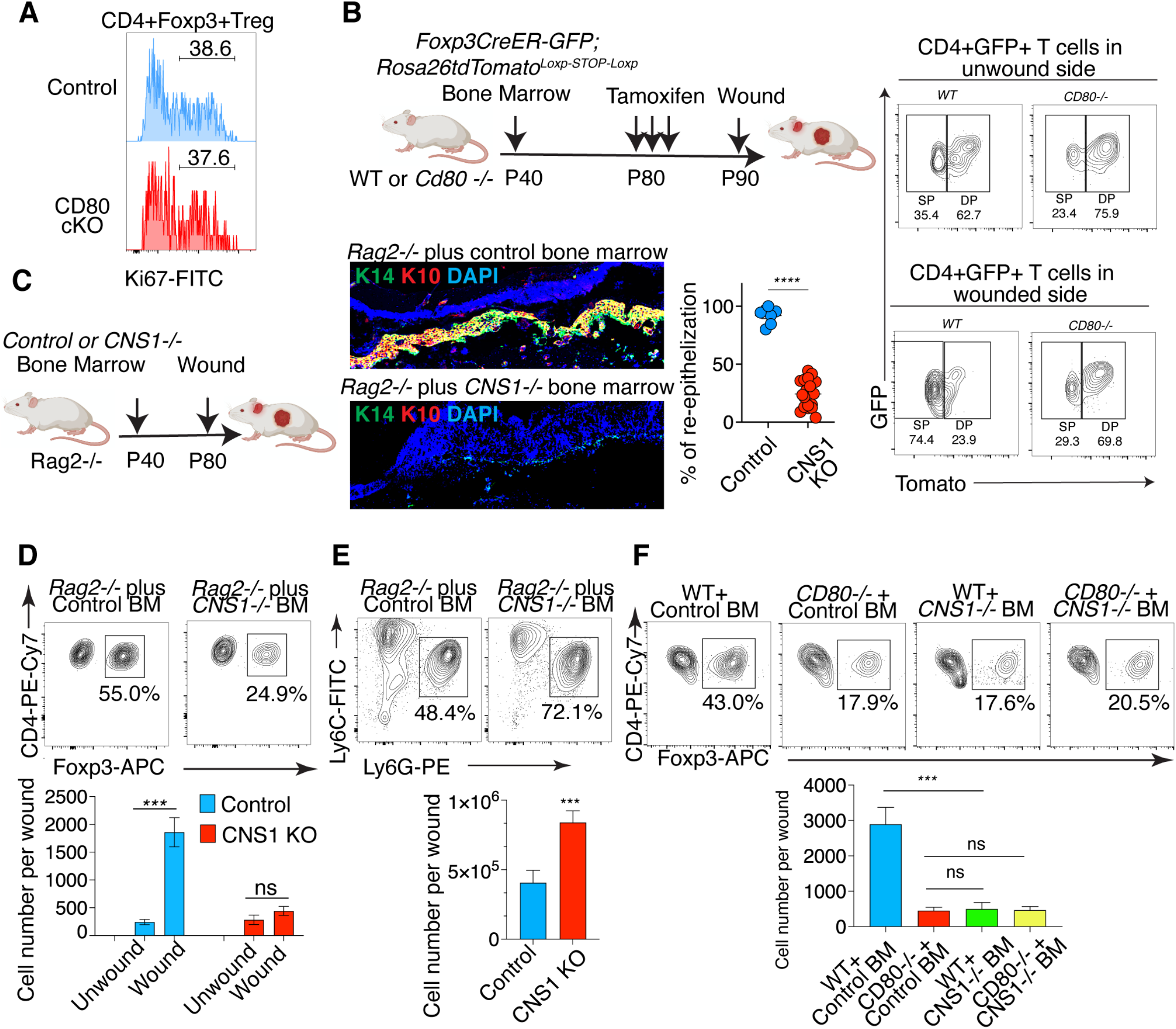
CD80 on HFSCs promotes de novo induction of Tregs during wound healing. **(A)** Flow cytometry quantification showing that proliferation (Ki67+) of Treg cells is comparable between WT and CD80 cKO wounds. **(B)** Experimental procedure and flow cytometry quantification of preexisting (DP: Tomato+GFP+) Tregs and de novo induced Tregs (SP: GFP+Tomato-) in unwounded or wounded WT or CD80 cKO mice. **(C)** Experimental procedure, IF image, and quantification of percentage of re-epithelialization (K14+) and number of differentiated cells (K10+) in Rag2-/- mice reconstituted with WT (control) or CNS1-/- (CNS1KO) bone marrow. **(D, E)** Flow cytometry quantification showing decrease of Tregs (**D**) and increase of neutrophils (**E**) in the CNS1-/- BM reconstituted mice, similar to CD80 BMC and CD80 cKO mice. **(F)** Flow cytometry quantification of Tregs in irradiated WT or CD80 KO mice reconstituted with control or CNS1-/- bone marrow.

Extrathymic induction of Treg cells from CD4+ T cells is dependent on the Foxp3 enhancer CNS1, which is dispensable for thymic Treg generation (Zheng et al., 2010). Interestingly, CNS1KO wounds had similar defects in re-epithelialization and differentiation, resembling those seen in Treg-depleted and CD80KO mice (Figure 4C). Immune landscape profiling of wounded CNS1KO mice found that CNS1KO wounds were significantly depleted of Tregs (Figure 4D), but were packed with neutrophils compared to control wounds (Figure 4E). This striking finding confirmed our hypothesis that extrathymic Treg cells generated *de novo* are the main population responsible for protecting HFSCs during wound healing. Next, we constructed bone marrow chimeras to further examine whether these *de novo*-generated Treg cells in the wound were induced through the action of HFSC-CD80. As predicted, the wounds of CNS1 KO and CD80 cKO mice were deficient in Tregs to a similar degree. Reconstitution of CD80 KO mice with CNS1 KO bone marrow did not result in any additional reduction of Treg cells when compared to CNS1 KO or CD80 cKO alone (Figure 4F). Taken together with our fate mapping experiment and genetic analysis, we conclude that HFSCs employ CD80 to induce *de novo* generation of Treg cells, and that these suppressive immune cells build a temporary niche, protecting HFSCs during wound repair.

### MHCII is not required for HFSC-mediated peripheral Treg induction

In the peripheral lymphoid organs, induction of Tregs is dependent on antigen presentation and concomitant co-stimulation by tolerogenic antigen presenting cells (Li and Rudensky, 2016). Intriguingly, a subset of murine keratinocytes were shown to express class II MHC (MHCII) (Tamoutounour et al., 2019). Analysis of the chromatin landscape of HFSCs revealed that the promoter of several MHCII genes are indeed accessible in HFSCs, even before wounding (Figure 5A). Flow cytometry and Imagestream examining the expression of MHCII further confirmed that HFSCs express MHCII before wounding, and that their expression level is elevated during wounding (Figure 5B, C).

**Figure 5.**
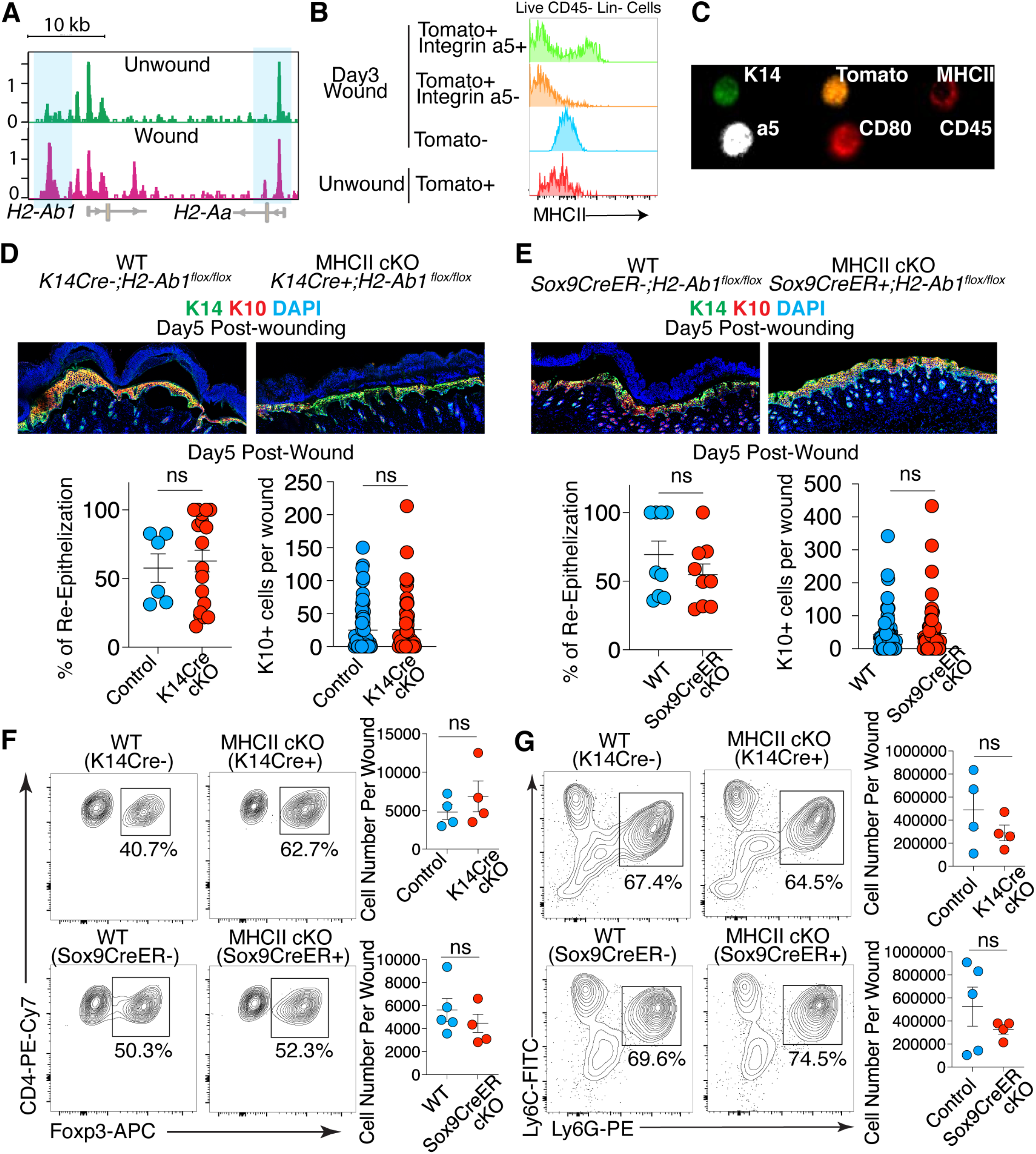
MHCII is dispensable for HFSCs to heal cutaneous wounds. **(A)** ATAC-seq peaks showing activation of class II MHC genes ( e.g. H2-Aa, H2-Ab1) in HFSCs, even before wounding. **(B, C)** Flow cytometry (**B**) and Imagestream (**C**) confirming elevated expression of MHCII on CD45-Integrin a5+ HFSCs in wounded skin. **(D, E)** IF and quantification of percentage of re-epithelialization (K14+) and number of differentiated cells (K10+) in WT or MHCII cKO driven by K14Cre (**D**) or Sox9CreER (**E**). **(F, G)** Flow cytometry quantification showing similar frequency and number of Tregs (**F**) and neutrophils (**G**) in control versus MHCII cKO mice.

To test the functional significance of SC-MHCII during wound healing, we knocked out class II MHC in the basal keratinocyte layer in the *K14-Cre; MHCII^flox/flox^* mice (MHCII cKO). To our surprise, we found no gross phenotypic differences in re-epithelialization or keratinocyte differentiation kinetics between MHCII cKO and Cre negative control mice (Figure 5D). One caveat to this model is that Keratin14 is also expressed in the thymic epithelium (Tamoutounour et al., 2019), which plays a crucial role in the maturation and selection of T lymphocytes during immune system development. Thus, to counteract any developmental defects in CD4 T lymphocytes, both Cre negative control and MHCII cKO mice were surgically grafted at 3 weeks of age with WT thymus into the donor renal capsule to supplant the endogenous lymphocytes. Upon reaching adulthood, these mice were wounded. Like their non-grafted counterparts, the MHC II cKO mice still successfully healed the wound (Supplementary Figure 5). To further test the importance of MHCII for wound repair, we used a third parallel model to postnatally target MHCII in HFSCs using a Tamoxifen-inducible Cre recombinase (*Sox9CreER; MHCII^flox/flox^*). In this model, Tamoxifen was given to 8 week-old mice immediately prior to wounding to activate Cre and ablate MHCII specifically in Sox9+ HFSCs of adult mice, such that loss of MHCII would not affect T cell development and repertoire. Like our previous models targeting MHCII in the skin, punctual ablation of MHCII in Sox9+ HFSCs in adult mice did not have an appreciable effect on the wound healing phenotype (Figure 5E). Additionally, the immune cell compositions in the wound beds, including the Treg and neutrophils populations, were comparable between WT and MHCII cKO mice in both models (Figure 5F, G). From these data we concluded that although MHCII is expressed by HFSCs, it is dispensable for stem cell-mediated Treg induction and wound repair.

### HFSCs provide an augment of co-stimulation signals to CD4+ effector T cells to induce Treg generation

Since MHCII on HFSCs did not seem to be crucial for priming Treg precursors in the wound, one possibility was that naïve CD4 T cells were being primed in the skin-draining lymph nodes, becoming activated, and then migrating into the wound bed. Consistent with this notion, coculture between HFSCs and naïve CD4 T cells in the presence of TGFβ did not result in Treg cell generation (Supplementary Figure 5B). Thus, we hypothesized that HFSCs may be affecting the effector CD4 T cells (Teff) in the wound. CD62L^Low^CD44^Hi^ CD4+ T_eff_ cells significantly increased as early as day 1 post-wounding in the wound-draining lymph nodes of both WT and CD80cKO mice (Figure 6A). By day 3, in WT mice, these non-Treg effector cells migrated into the wound, were highly proliferative, and produced a large amount of IL2 (Figure 6B, C). In contrast, proliferation and IL-2 production of Teff cells were significantly blunted in wounded CD80cKO mice (Figure 6B, C).

**Figure 6.**
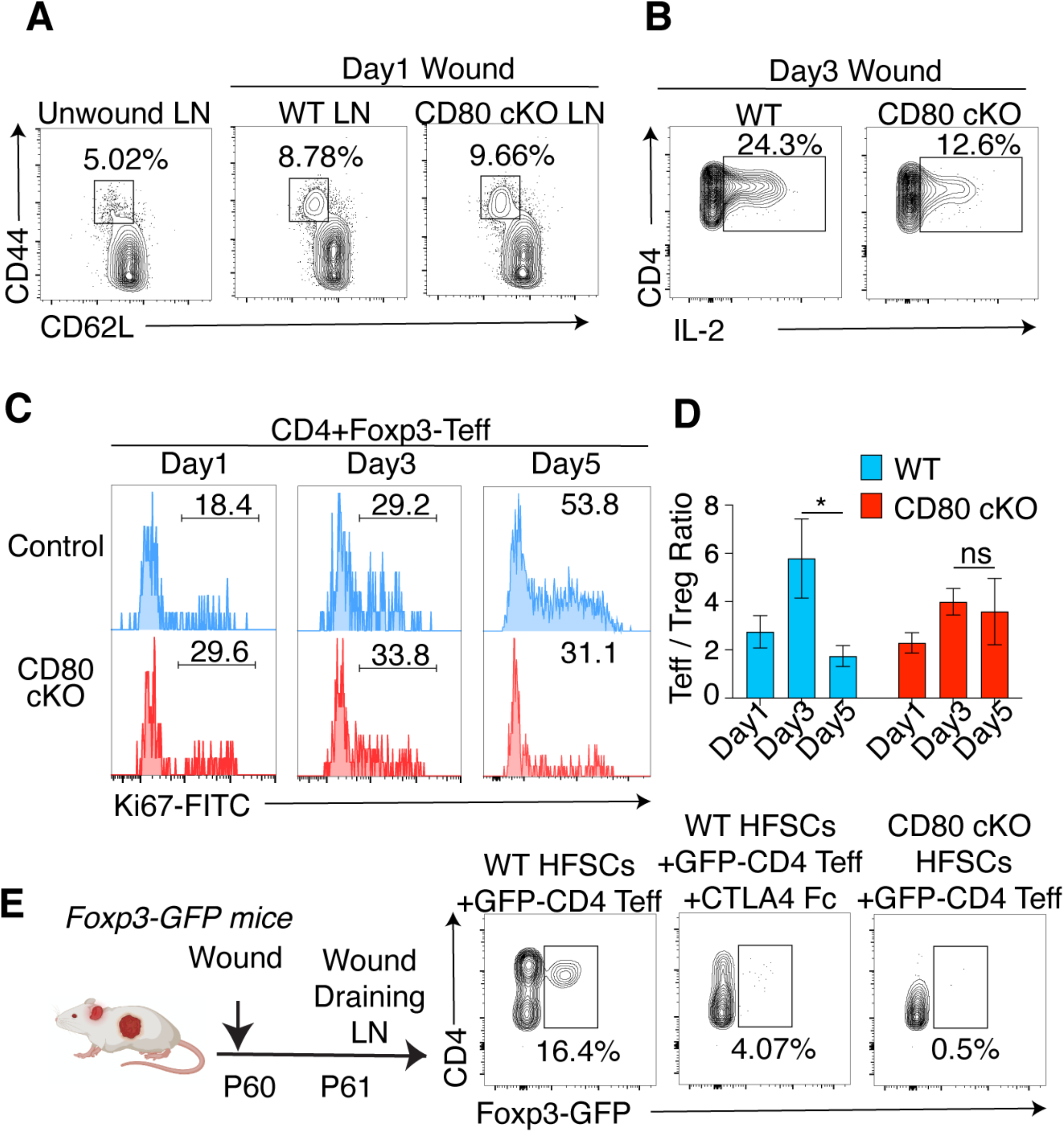
HFSCs induce wound-infiltrating CD4 effector T cells to differentiate into Treg cells during wound repair. **(A)** Flow cytometry quantification showing similar increases of CD62L^low^ CD44^Hi^ CD4 effector T cells (Teff) in the wound draining lymph nodes (LN) in WT and CD80 cKO mice at day 1 post-wound. **(B, C)** Flow cytometry quantification showing decreased IL2 production **(B)** and Ki67+ proliferative Teff cells **(C)** in CD80 cKO wounds compared to WT wounds. **(D)** Teff/Treg ratio in WT versus CD80 cKO wounds showing defective conversion of Teff cells into Treg cells in CD80 cKO wounds. **(E)** GFP-CD44^Hi^CD62^Low^CD4 Teff cells isolated from the LNs of the wounded Foxp3-GFP reporter mice were co-cultured with WT or CD80 cKO HFSCs or WT HFSCs incubated with CTLA4 Fc protein, in the presence of IL2 and TGFβ. Note that only WT HFSCs with functional CD80 were able to induce Teff cells to differentiate into Foxp3+ Treg cells.

These data led us to hypothesize that HFSCs, rather than priming naïve CD4 T cells, provide a boost of co-stimulation signals to effector CD4 T cells, triggering their differentiation into Treg cells in an environment enriched for TGFβ, such as in the wound bed (). Consistent with this hypothesis, we measured the Teff/Treg ratio in both WT and CD80 cKO mice. While this ratio increased in WT wounds at day3 post-wounding, it decreased in the days afterwards, presumably due to the differentiation of the infiltrating Teff into Treg cells. In CD80 cKO mice, while we observed an increased Teff/Treg ratio at day3, this ratio was maintained thereafter (Figure 6D). To further test this hypothesis, we established a co-culture assay. We first wounded Treg reporter mice (*Foxp3-GFP*) and isolated GFP^neg^CD62L^Low^CD44^Hi^ CD4 Teff cells from the wound-draining lymph nodes (Figure 6E). These Teff cells were co-cultured with HFSCs isolated from WT or CD80KO mice. In parallel, we isolated GFP-Teff cells and cocultured them with WT HFSCs that were coated with CTLA4 Fc, which blocks SC-CD80 function. After incubation for 5 days, co-culture with WT HFSCs promoted activation of Foxp3 in a significant number of CD4 Teff cells. On the other hand, Treg cells were absent after coculturing with WT HFSCs coated with CTLA4 Fc, or with CD80 KO HFSCs (Figure 6E).

### Intimate interactions between HFSCs and T cells are vital for wound healing

Our study has revealed an intimate dialogue between HFSCs and CD4 T cells which shape a Treg-mediated immune-privileged temporary niche. This niche is critical for wound-healing HFSCs to evade the inflammation caused by wound-infiltrating neutrophils. However, it is unclear why HFSCs would have to activate a co-stimulation ligand such as CD80 to induce *de novo* Treg generation rather than secreting cytokines to directly recruit natural Treg cells from the lymphoid organs into the wound. One possibility is that by engaging with Teff through CD80/CD28 interactions, HFSCs can induce Treg generation in close proximity, and in doing so, provide close and precise protection. On the other hand, if Tregs are recruited by cytokines, localization of Treg cells directly adjacent to HFSCs may prove more difficult. To test this scenario, we treated CD80 cKO mice with a CD28 superagonist (SA) antibody which is known to induce proliferation and expansion of Treg cells. As we expected, subcutaneous injection of CD28 SA antibody into the wound bed of CD80 cKO mice dramatically increased Treg numbers in the wound (Figure 7A). In fact, the Treg number was rescued to a level similar to that produced by WT mice during wounding (Figure 7B). As a result, neutrophils were reduced by CD28 SA treatment. However, even though Treg number was rescued to a WT level in the wound, phenotypically, the wound healing defects of CD80 cKO mice could not be rescued by CD28 SA antibody treatment (Figure 7C). Remarkably, IF imaging showed that Treg cell numbers were significantly increased in wounded skin after CD28 SA antibody treatment, but Treg cells remained absent in the wound bed of the CD80 cKO mice (Figure 7D). Consistent with our hypothesis, even though the neutrophil number was reduced in the CD28 SA antibody-treated wound, without localization of Tregs to the HFSCs, the remaining neutrophils still infiltrated the wound bed, hindering the re-epithelization process.

**Figure 7.**
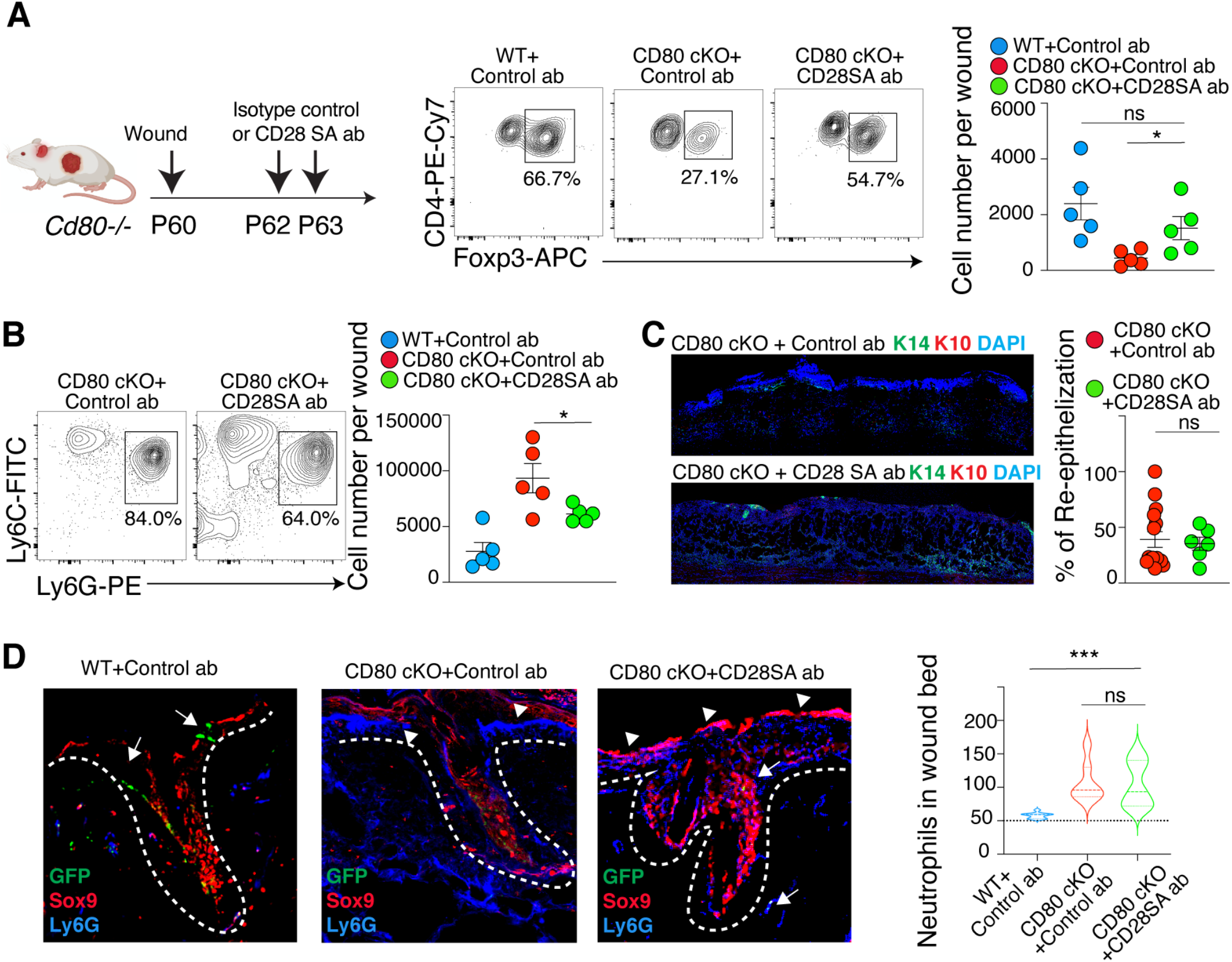
Intimate crosstalk between HFSCs and CD4 T cells is necessary to prevent collateral damage from neutrophils during wound repair. **(A, B)** Experimental procedure and flow cytometry quantification of Treg cells (**A**) and neutrophils (**B**) after subcutaneous injection of CD28 super-agonist (SA) antibody. **(C)** IF and quantification of percentage of re-epithelialization (K14+) in CD80 cKO mice receiving isotype control or CD28 SA antibody treatment. **(D)** IF staining and quantification showing that accumulation of Tregs within the wound bed prevents localization of neutrophils to Sox9+ cells in WT mice (left). In contrast, in CD80 cKO mice, neutrophils accumulate in the wound bed (mid), and CD28 SA antibody treatment cannot rescue the neutrophil accumulation (right).

Collectively, we conclude that HFSCs, after migration into the wound bed, acquire an immune modulatory capacity which allows them to directly engage with infiltrating effector CD4 T cells and induces their to differentiation into Treg cells. Through this vital interaction, these *de novo*-generated Treg cells build a close barrier surrounding the HFSCs and shut down local neutrophil-mediated inflammation, allowing HFSCs to proliferate and differentiate to repair a wound.

## Discussion

HFSCs comprise a vital cellular reservoir for regenerating the skin, yet their location within a barrier tissue constantly exposes them to various inflammatory insults. In homeostasis, they reside within the protective, immunosuppressive hair follicle niche, but during cutaneous wounding, this niche is not only disrupted, but also HFSCs must migrate out of their niche and enter into the highly inflammatory environment of the wound bed to repair the wound. In this study, we unveiled a remarkable molecular feature of HFSCs that helps stem cells endure inflammation after they migrate into the wound bed. In order to achieve this feat, HFSCs upregulate CD80, a cell-surface marker canonically expressed on antigen-presenting cells. Surprisingly, unlike tolerogenic antigen-presenting cells, HFSCs expressing CD80 are able to modulate the strength of the co-stimulation signals delivered to effector CD4 T cells that infiltrate into the wound. In a wound microenvironment enriched with TGFβ, this augmentation of the costimulation signal triggers Teff cells to differentiate into Treg cells which, in turn, prevent accumulation of pro-inflammatory innate immune cells around HFSCs, allowing HFSCs to generate new epidermis.

### Immune modulatory capacity of adult stem cells

It has long been postulated that adult SCs are a group of vulnerable cells that need to be protected within an immune privileged niche. Our study finds that adult stem cells are actually an integral part of the immune modulatory network in the body. This represents a major paradigm shift in our understanding of how SCs can drive immune responses and tissue regeneration. It has been recently reported that intestinal and epidermal SCs express class II MHC, and that presence of MHCII can increase cytokine production from tissue-resident CD4 T cells which, in turn, promotes SC functions under homeostatic conditions (Biton et al., 2018; Tamoutounour et al., 2019). Interestingly, upon wounding, when HFSCs migrate out of their natural niche and face robust inflammation on their own, these cells can activate immune modulatory program and express both class II MHC and a costimulatory molecule. This finding raises the tantalizing possibility that HFSCs under inflammatory conditions may mimic immune cells and act as non-professional antigen presenting cells. To our surprise, in contrast to under homeostatic conditions, we find that MHCII is actually dispensable during wound repair. It appears only the CD80-mediated co-stimulatory signal is required to actively sculpt a temporary protective niche comprised of *de novo*-generated Treg cells. This may be due to the lack of necessary tertiary lymphoid structures in the wound bed to achieve optimal activation of naïve T cells which is required for iTreg induction. Therefore, HFSCs have chosen the ideal mechanism by instead modulating effector T cells, by providing a boost of co-stimulatory signals to Teff cells to induce Treg generation.

Though peripheral Treg induction has previously been attributed to signals derived from antigen presenting cells, we present a novel pathway in which epithelial SCs are able to co-opt canonically immune programs in response to inflammation to induce Treg generation. Strikingly, we further demonstrated that, by initiating this intimate interaction with T cells, Treg cells can be generated in close proximity to the wound-healing HFSCs. This interaction may help newly-generated Treg cells precisely pinpoint the location of HFSCs in order to suppress local inflammation and effectively protect the most critical group of cells during skin regeneration.

This newfound appreciation of immune modulatory capacity of adult tissue SC also raises major questions about the role that SCs play in the modulation of inflammatory diseases. Future studies are needed to investigate whether impaired SC function may underlie the intractable skin inflammation displayed by patients with inflammatory dermatoses such as atopic dermatitis or alopecia areata, particularly in those who have impaired regulatory T cell responses. Another question that arises from this study is whether inability to turn on immunomodulatory programs in SCs is responsible for the lack of wound healing in chronic wounds, and whether these programs can be targeted therapeutically early in wound development to prevent the self-perpetuating cycle of barrier compromise and inflammation that prevents many chronic wounds from healing.

The new findings from this study have particularly important implications in cancer. Recently, it has been revealed that a group of TGFβ-responding tumor initiating stem cells have the unique capacity to evade immune surveillance and survive robust immunotherapy treatment (Miao et al. 2019). Interestingly, these cancer stem cells also acquire CD80, which was shown to directly modulate cytotoxic T cell activities. Driven by the observations from the current study, it will be critical for future research to investigate how cancer stem cells orchestrate Treg responses during cancer progression, and how cancer stem cells hijack the stem cell-specific immune modulatory features that are activated during wound healing.

Overall, this study points to an intricate stem cell-driven orchestration of immunity during wound repair that has major implications for the way we think about pathogenic immune responses.

### The role of T cells in cutaneous wound repair

The immune responses evoked during cutaneous wound repair have always been considered to be dominated by innate immunity, and the roles of adaptive immune cells have long been overlooked. This study has revealed the key role of CD4 T cells in driving wound healing. Most CD4+ T cells initially infiltrating a cutaneous wound express an effector T cell phenotype, suggesting that they are activated in the lymphoid organs. Interestingly, these effector CD4 T cells begin to increase in the skin-draining lymph nodes as early as day one, and begin to infiltrate the wound. It is intriguing to see a T cell response at this early time point. In the future, it will be important to determine the antigen specificity of these wound-infiltrating CD4 T cells, especially the clones that are induced to become Treg cells. It is also possible that these T cells are primed even before wounding. Future studies are needed to examine the role of various factors, such as skin microbiota, in priming these T cell clones in the first place--in particular, whether these primed T cells are a result of the early-life colonization by microbiota.

Our study also provided an important piece of evidence supporting the vital role of Treg cells during tissue regeneration and wound healing. During cutaneous wound repair, Tregs have been shown to attenuate IFNγ production and proinflammatory macrophage accumulation (Josefowicz et al., 2012). Here, we demonstrate that Treg cells can provide close protection to the wound healing stem cells by shielding them from the damage inflicted by inflammatory neutrophils. In line with this conclusion, recent studies have shown that Treg cells can blunt IL17 production which is vital for neutrophil recruitment and accumulation in the wound (Mathur et al., 2019). Interestingly, our neutrophil depletion experiment only partially rescued wound re-epithelization and differentiation. This data suggests that preventing neutrophil accumulation might represent only a portion of the contributions of Tregs to cutaneous wound repair. In the context of muscle wound repair, Tregs have also been shown to secrete a variety of EGFR ligands, such as Amphiregulin (Arpaia et al., 2015). Whether these processes play a role in facilitating HFSC proliferation and differentiation still await future study to determine.

---

**Supplementary Figure 1.**
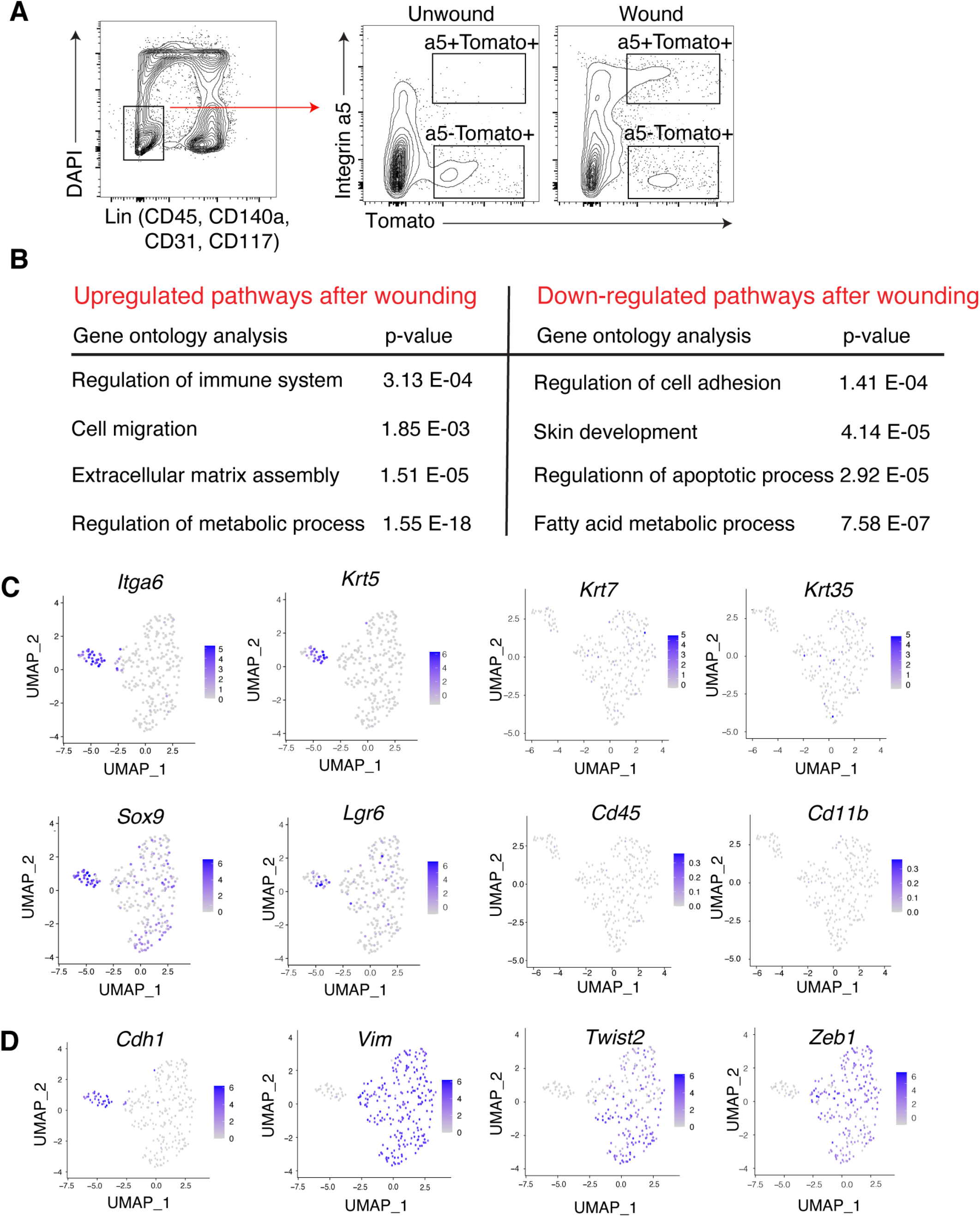
**(A)** Sorting strategy to isolate HFSCs from unwounded or wounded skin **(B)** GO term analysis of pathways upregulated/downregulated in HFSCs after wounding **(C, D)** t-SNE plots showing the signatures that are enriched in Integrin a5 hi migratory HFSCs after wounding

**Supplementary Figure 2.**
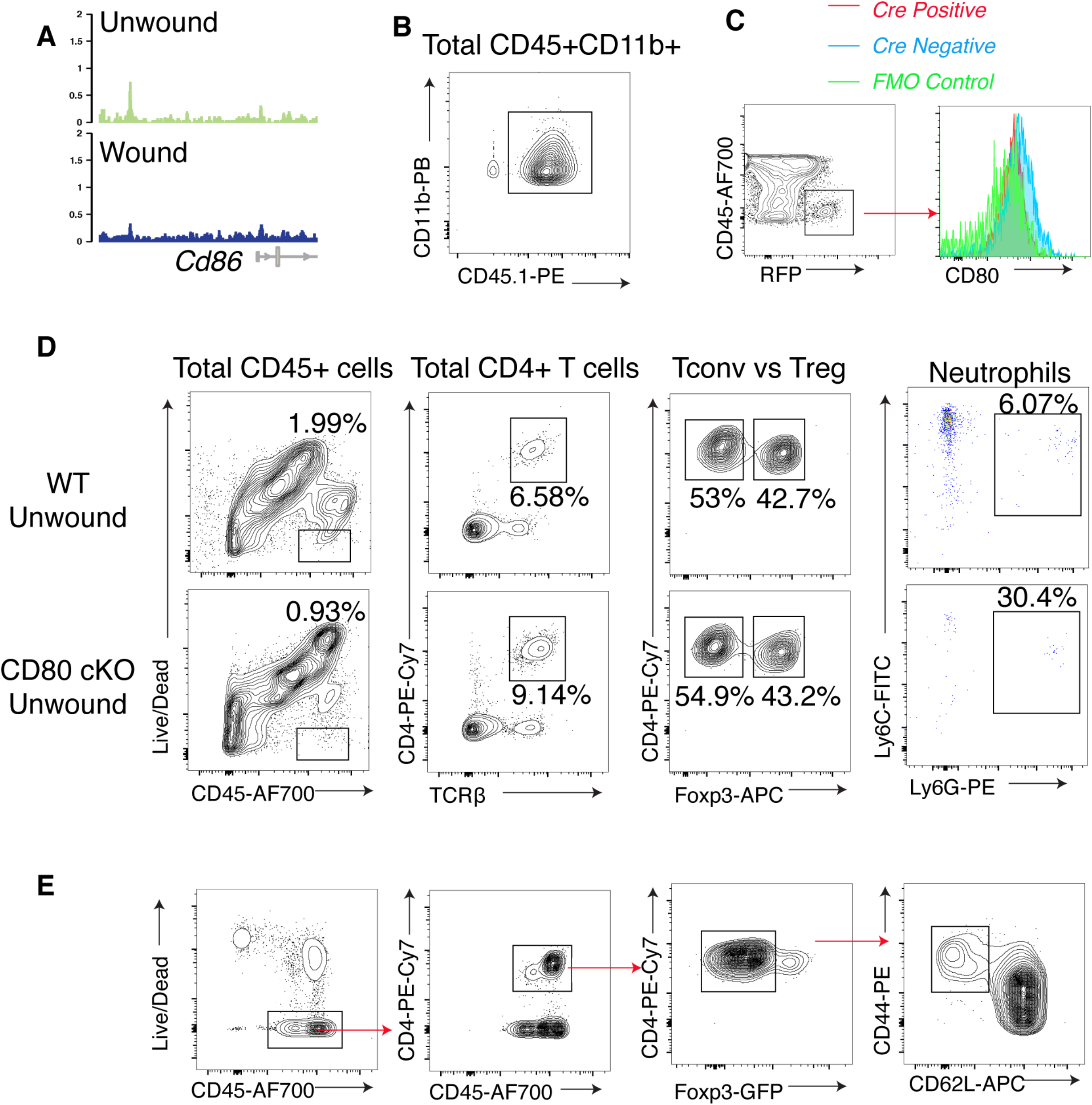
**(A)** ATAC-seq peaks showing Cd86 is not activated in HFSCs **(B)** Flow cytometry quantification showing that the majority of CD11b+ myeloid cells are derived from CD45.1+ donor bone marrow **(C)** Flow cytometry quantification showing efficient silencing of CD80 after in utero injection of lentivirus carrying a sgRNA targeting *Cd80*. **(D)** Flow cytometry quantification showing that the major immune populations (e.g Treg and neutrophils) are similar in unwounded skin between WT and CD80 cKO mice. **(E)** Sorting strategy for isolating CD4 effector T cells for co-culture.

## EXPERIMENTAL MODEL AND SUBJECT DETAILS

### CONTACT FOR REAGENT AND RESOURCE SHARING

Further information and requests for reagents and resources should be directed to the Lead Contact, Yuxuan Miao or Elaine Fuchs.

#### Animals

Sox9CreER and K14Cre (Fuchs Lab) have been previously described and were backcrossed to C57/Bl6 background for ten generations. *Foxp3^GFP-CreER^* and *Foxp3^ΔCNS1-gfp^* (CNS1-KO) were obtained from the Rudensky Lab at Memorial Sloan Kettering Cancer Center. Wild-type C57BL/6J, B6;129S6-Gt(ROSA)^26Sortm9(CAG-tdTomato)Hze^/J (*R26-LSL-tdTomato*), B6.129S4-*Cd80^tm1Shr^*/J (*CD80^-/-^*), B6;129-Gt(ROSA) 26Sor^tm1(CAG-cas9*,-EGFP)Fezh^/J (*R26-LSL-Cas9*), B6.129(Cg)-*FOXP3^tm3(DTR/GFP)Ayr^*/J (*Foxp3-DTR*), B6(Cf)-*Rag2^tm1.1Gn^*/J (*RAG2^-/-^*), B6.SJL-Ptprc^a^ Pepc^b^/BoyJ (*CD45.1*), and *MHCIIfl/fl (B6.129X1-H2-Ab1 ^tm1Koni^/J)* mice were obtained from The Jackson Laboratory. Tregs were depleted in Foxp3-DTR mice using a modified protocol (Green et al., 2017). Briefly, diphtheria toxin (DT, Sigma-Aldrich) was diluted to 2.5 ug/ml, and 200 ul (0.5 ug) was injected intraperitoneally (i.p.) into WT or Foxp3-DTR mice at Day −7, −3, and 0. On day 0 mice were wounded. Neutrophils were depleted in CD80 cKO mice via i.p. injection with 50ug CXCL5 monoclonal neutralizing antibody (R&D systems; mAb 433) at day 0, 1, and 3 post-wound. To replete Tregs in the wound, 100 ug CD28 superagonist antibody (clone D665) was i.p. injected into CD80cKO mice at day 0, 1, and 3 post-wound. For treatment of mice with tamoxifen, daily i.p. injection of 100 μg for five consecutive days was performed starting at 7 days prior of wounding. All mice were maintained in an Association for Assessment and Accreditation of Laboratory Animal Care (AAALAC)-accredited animal facility, and procedures were performed using IACUC-approved protocols that adhere to NIH standards. The animals were maintained and bred under specific-pathogen free conditions at Comparative Bioscience Center at Rockefeller University and all the procedures are in accordance with the Guide of the Care and Use of Laboratory Animals.

#### Cell Lines

HEK293T cells were used for packaging lentivirus and were cultured in Dulbecco’s modified eagle medium (DMEM) with 10% FBS, 100 units/mL streptomycin, 100mg/mL penicillin, and 2mM glutamine.

### METHOD DETAILS

#### *In utero* lentiviral transduction

*Cd80* was excised specifically in the skin of *K14Cre; Rosa26-LSL-Cas9* mice via the ultrasound-guided *in utero* injection of the lentivirus carrying *Cd80*sgRNA. Briefly, at 9.5 days gestation (E9.5), when the embryo consists of a single layer of progenitors, female mice were anesthetized with isoflurane (Hospira) and low-titer lentivirus was injected into their amniotic sacs in order to selectively transduce a small number of progenitors within the surface ectoderm that eventually develop into the skin (Beronja et al., 2010). Once the virus is integrated (~24 hr post-infection), the DNA carried by the lentivirus is stably propagated to the progeny of the epithelial progenitors (Beronja et al., 2010).

#### Bone marrow chimera construction

Recipient mice were lethally irradiated (700 rads), which ensures depletion of endogenous hematopoietic stem cells. Radiation was delivered in two equal doses (350 rads each), 4 hrs apart, to minimize damage to gastrointestinal and pulmonary cells. Bone marrow cells were isolated from donor femurs and 1 x 10^7^ bone marrow cells were adoptively transferred through tail vein injection into irradiated recipient mice. Animals were maintained in autoclaved cages with autoclaved water and fed sulfatrim antibiotic-containing food. Six to eight weeks post-transplantation, recipient mice were used for experiments.

#### Partial-thickness wound model

Mice were wounded in telogen (postnatal day P60, or postnatal day P120 for bone marrow chimeras). Briefly, mice were shaved, hair was removed using depilatory cream, and a Dremel drill head was used to gently scrape the back skin of anesthetized mice. In order to standardize the wound depth, the number of touches performed with the Dremel drill was determined by inspecting the wounded skin for the first signs of erythema and pinpoint bleeding. This method removes the epidermis and upper HF, including most infundibulum and isthmus cells, but leaves the HF bulge intact.

#### Fluorescence Activated Cell Sorting and Flow Cytometry

To isolate HFSCs for *in vitro* co-culture, the back skins from wounded WT or *Cd80-/-* mice were dissected and scraped with a blunt blade from the dermal side to remove fat. Skin was placed with the dermal side down in trypsin for digestion for 30 minutes. A single cell suspension was then prepared and stained with an antibody cocktail. Integrin a6+CD34+Sca1-HFSCs were sorted and cultured in Y medium with calcium concentration of 650 um. For immune profiling of the wound, the wound was placed in cold PBS for 15 minutes to remove the scab. The remainder of the wounded tissue was minced in digestion media [RPMI (Thermo Fisher) with HEPES (1:40, Thermo Fisher), MEM 100X (1:100, Thermo Fisher), Na pyruvate (1:100, Thermo Fisher), penicillin (100ug/ml), streptomycin (100 units/ml), gentamicin (1:500, Thermo Fisher), ß-mercaptoethanol (1:1000)]. The tissue was digested with in Liberase (25 μg/ml) (Roche) for 120 minutes at 37 °C with an adapted protocol (Naik et al., 2017). Single cell suspensions were obtained and cells were stained using an antibody cocktail prepared at predetermined concentrations in a staining buffer (PBS with 5% FBS and 1% HEPES). Fixable Live/Dead was used to exclude dead cells. Stained cells were analyzed with LSR II Analyzer (BD Biosciences) and with the FlowJo program. To sort effector CD4 T cells for *in vitro* co-culture, the wound-draining lymph nodes were collected at day 1 post-wound using Foxp3-GFP mice. CD44^Hi^ CD62L^low^ CD4 effector T cells were sorted to be co-cultured with WT or CD80 KO HFSCs at a 1:1 ratio and incubated in T cell medium for five days in the presence of recombinant IL2 and TGFβ.

#### Single cell RNA-sequencing

Single cell RNA-seq was performed using the Smart-seq2 protocol with modifications (Picelli et al., 2014; Yang et al., 2017). Briefly, single cells from wounded and unwounded skin were sorted into 96-well plates containing 2ul lysis buffer [0.2%Triton X-100, 20U/uL SUPERase In RNAse inhibitor (Thermo Fisher), 0.25 uM oligo-dT30VN primer, and 1:8,000,000 external RNA (ERCC) spike-in RNAs control (Ambion)]. Samples were then flash frozen with liquid nitrogen and stored at −80C until processing. The plates were thawed on ice and then heated at 72C for 3 min to lyse the cells. Reverse transcription was then performed using 4U/ml Maxima H-transcriptase (Thermo Fisher), utilizing the template switching reaction and PCR pre-amplification (15 cycles) according to the protocol (Picelli et al., 2014). The cDNA PCR products were cleaned and purified with AMPure XP beads. To evaluate for and exclude low quality libraries or empty wells, RT-qPCR was used to amplify the housekeeping gene actin. Illumina sequencing libraries were then prepared using the Nextera XT DNA library preparation kits (Illumina). Samples were then pooled and sequenced with the Illumina Nextseq 500 machine using a 75-bp paired-end-read setting.

#### Immunofluorescence and Imaging

Wounded and unwounded skin was fixed in 4% paraformaldehyde immediately after dissection for 1hr at 4C and washed three times in PBS. The tissue was incubated in 30% sucrose in PBS at 4C overnight. Tissue was then embedded in OCT (Tissue Tek), frozen, and sectioned (12-20um). Cryosections were permeabilized, blocked, and stained with the following primary antibodies: K10 (Rabbit, 1:1000, Fuchs Lab), K14 (Chicken, 1:1000, Fuchs Lab), mCD80 (Goat, 1:100, R&D), GFP (Chicken, 1:1000, Abcam), RFP (Rabbit, 1:1000, Ly6G (Rat, 1:100, Biolegend), integrin a5 (Rat, 1:100, BD). The slides were then stained with secondary antibodies conjugated with Alexa 488, 546, or 647 (Life Technologies) and imaged on Zeiss Axio Observer Z1 equipped with ApoTome.2 or Leica Stellars 8 Laser Scanning Confocal microscope. The images were collected using Zeiss ZEN software and analyzed using Fiji/ImageJ software.

#### Statistics

Statistical analysis for microscopy quantifications was performed in Prism 9 (GraphPad). Column data was first analyzed using D’Agostino and Pearson normality testing. For data not normally distributed, the two-tailed Mann-Whitney test was performed with a 95% confidence interval. For normally distributed data, the unpaired two-tailed student’s t test was performed with a 95% confidence interval. Data are presented as mean + SEM. Significant differences between groups were noted by asterisks (* p<0.01; ** p<0.005, *** p<0.001, *** p<0.0001). Group size was determined by power analysis on the basis of preliminary experiment results. Experiments were performed unblinded.

